# Suppression of PIK3CA-driven epileptiform activity by acute pathway control

**DOI:** 10.1101/2021.03.03.433821

**Authors:** Achira Roy, Victor Z. Han, Angela M. Bard, Devin T. Wehle, Stephen E. P. Smith, Jan-Marino Ramirez, Franck Kalume, Kathleen J. Millen

**Affiliations:** Center for Integrative Brain Research, Seattle Children’s Research Institute, Seattle, Washington, USA; Departments of Biology University of Washington, Seattle, Washington, USA; Graduate Program in Neuroscience; Departments of Pediatrics University of Washington, Seattle, Washington, USA; Departments of Neurological Surgery University of Washington, Seattle, Washington, USA; Departments of Physiology and Biophysics University of Washington, Seattle, Washington, USA; Departments of Pharmacology, University of Washington, Seattle, Washington, USA

**Keywords:** PI3K, epilepsy, mouse model, preclinical drug screens, BKM120, RAD001, AZD5363, intractable, pyramidal neuron, hippocampus, electrophysiology

## Abstract

Patients harboring mutations in the PI3K-AKT-MTOR signaling pathway often develop a spectrum of neurodevelopmental disorders including epilepsy. A significant proportion of them remain unresponsive to conventional anti-seizure medications. Understanding mutation-specific pathophysiology is thus critical for molecularly targeted therapies. We previously determined that mouse models expressing patient-related activating mutation in *PIK3CA* are epileptic and acutely treatable with PI3K inhibition, irrespective of dysmorphology (Roy et al. 2015). Using the same mutant model, we have now identified physiological mechanisms underlying the dysregulated neuronal excitability and its acute attenuation. We show that Pik3ca-driven hyperexcitability in hippocampal pyramidal neurons is mediated by changes in multiple non-synaptic, cell-intrinsic properties. These are distinct from mechanisms driving epilepsy in TSC/RHEB models. Further, we report that acute inhibition of PI3K or AKT, but not MTOR, suppresses the intrinsic epileptiform nature of the mutant neurons. These data represent an important step towards precision therapeutics against intractable epilepsy, using pathway drugs originally developed as anti-cancer agents.

## INTRODUCTION

Mutations in the PI3K-AKT-MTOR signaling pathway, long studied for roles in cancer (Madsen 2020; Yang et al. 2019), also cause clinically important developmental brain overgrowth syndromes. Affected individuals display phenotypes ranging from dysplastic megalencephaly, hemimegalencephaly and focal cortical dysplasia, as well as comorbidities including hydrocephalus, autism and intellectual disability (Bast et al. 2006; Blumcke et al. 2011; Mirzaa 2018; Stafstrom and Carmant 2015; Dobyns and Mirzaa 2019; Crino 2016). These mutations also cause focal epilepsy, representing 25-50% of all cases of intractable (treatment-resistant) epilepsy in children (Bast et al. 2006; Blumcke et al. 2011; Kim and Lee 2019; Mirzaa 2018; Stafstrom and Carmant 2015). Traditionally, anti-seizure drugs targeting generic ion channels are identified by evoking acute seizures in wild-type rodents (Wilcox, West, and Metcalf 2020; Kehne et al. 2017; Barker-Haliski and White 2019), which do not mimic clinical epilepsies associated with specific genetic mutations and hence are largely ineffective. Since PI3K pathway mutations are now known to cause intractable epilepsy, repurposing pathway-targeted anti-cancer drugs offers a tantalizing opportunity to fundamentally shift the therapeutic approach towards intractable epilepsy. Presently, the MTOR inhibitor rapamycin and its analogs are the sole pathway-related drugs used clinically to treat epilepsy, primarily in patients with rare *TSC1/2* deletion mutations which enhance downstream MTOR signaling(Stafstrom 2019; Crino 2016; Kim and Lee 2019). Yet, these treatments remain effective only to a modest degree, suggesting that MTOR activation may not be the sole arm of this complex signaling pathway accounting for all PI3K pathway-driven epilepsies (Cho 2011; Meng et al. 2013; Roy et al. 2015).

Here we assess acute mechanisms driving epileptiform activity in *Nestin-cre;Pik3ca^E545K^* mouse model harboring a patient-related activating mutation in *Pik3ca*, encoding the p110a catalytic subunit of PI3K (Roy et al. 2015). We report that Pik3ca overactivation causes intrinsic neuronal changes leading to hyperexcitability. We further establish that this epileptiform activity is acutely dependent on AKT, but not MTOR, regulation. Our study defines key aspects of acute neuronal dysregulation in an important genetic epilepsy model, which we leverage to repurpose the available PI3K pathway inhibitors against intractable epilepsy.

## RESULTS

### *In vivo* recordings demonstrate network hyperexcitability in mutant CA1

*Pik3ca* mutant mouse models recapitulate patient phenotypes including brain overgrowth, cortical dysplasia, hydrocephalus and epilepsy, with phenotypic severity dependent on the mutant allele and its time of activation (Roy et al. 2019; Roy et al. 2015). Moreover, this developmental epilepsy is dissociable from dysmorphology and acutely suppressible by 1-hour *in vivo* administration of the pan-PI3K inhibitor BKM120 (Maira et al. 2012; Roy et al. 2015). To identify the underlying dynamic mechanisms, we performed *in vivo* and *in vitro* electrophysiological studies primarily focused on hippocampal pyramidal neurons, since the connection between epilepsy and hippocampal pathophysiology is well documented in patients and model systems, including ours (Roy, Millen, and Kapur 2020; Chatzikonstantinou 2014; Mazumder, Patial, and Singh 2019; Berdichevsky et al. 2013; Roy et al. 2015).

We investigated *in vivo* changes in the hippocampal CA1 network activity of ∼P70 *Nestin-cre;Pik3ca^E545K^* mutant mice relative to control littermates using local field potential (LFP) recordings, while simultaneously monitoring cortical surface activity by electrocorticography (ECoG). These recordings showed spontaneous interictal spike activity either restricted to the cortex or the hippocampus exclusively, or concurrent to the two regions in the mutant mice (Figure 1A-C; Figure 1 – figure supplement 1A-D). A broad spectrum of spike patterns was observed, consisting of single or groups of interictal spikes, the simplest identifiable unit of epileptiform activity (McCormick and Contreras 2001), as well as trains of sharp spikes, low-frequency slow waves or high-frequency low-amplitude “brushing events”. These epileptiform events were observed exclusively in the mutant mice. Within the mutant group, the event frequency was significantly higher in the hippocampus than in the neocortex (Figure 1D). Power spectrum analyses revealed that the mutant hippocampi exhibit significantly higher power in the gamma frequency bands (Figure 1E-H), which is often indicative of seizure onset (Hughes 2008; Lee, Spencer, and Spencer 2000). These observations indicate that Pik3ca overactivation caused significant neural hyperexcitability in our mice, predominantly in the hippocampus.

**Figure 1:**
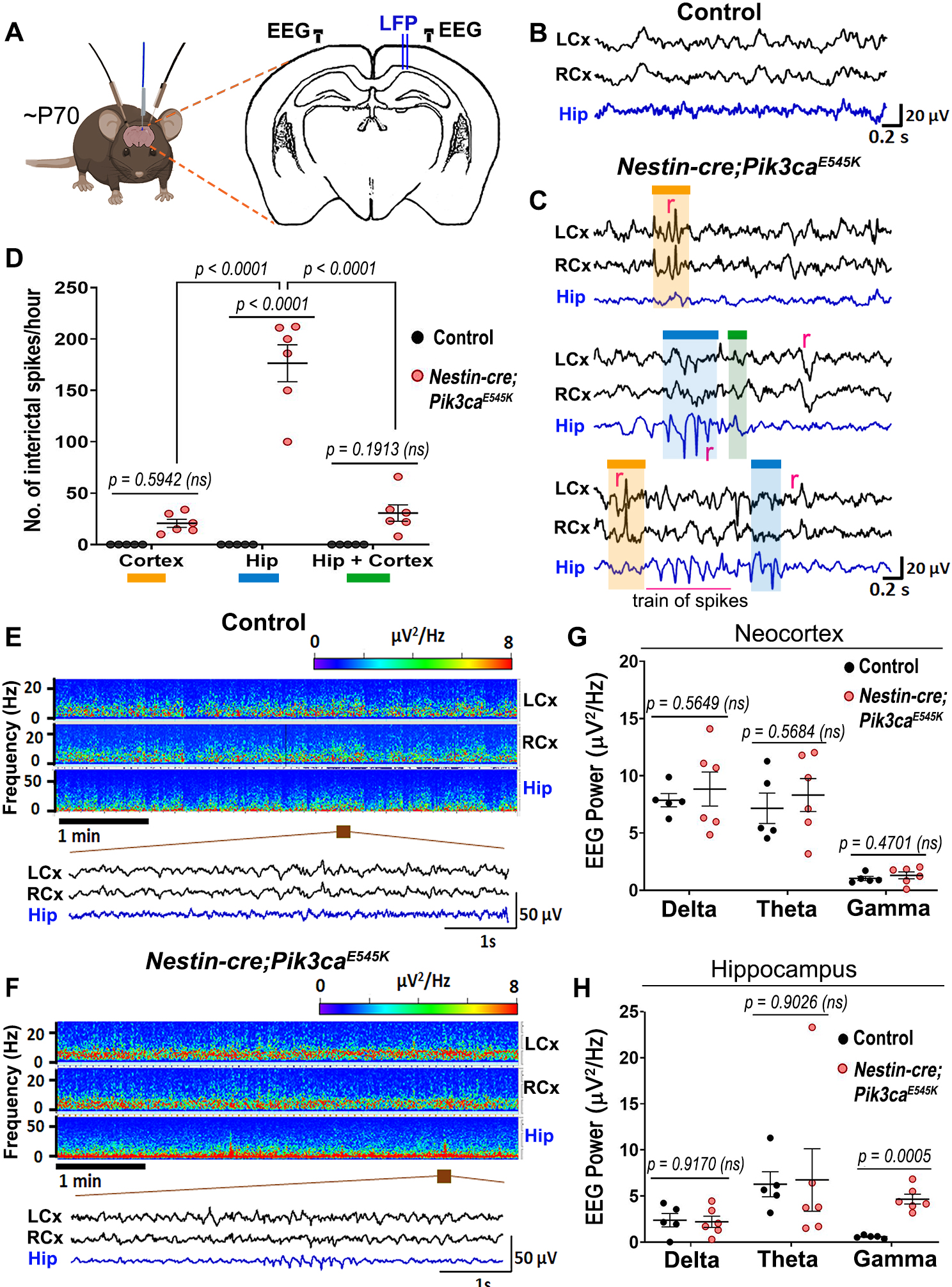
*Nestin-cre;Pik3ca^E545K^* mutant brains show higher neural excitability. (**A**) Schematic shows electrode placement for EEG-ECoG recordings in ∼P70 *Nestin-cre;Pik3ca^E545K^* and control littermates. LFP, local field potential. (**B,C**) Compared to controls, the mutants showed significantly higher frequency of regional (r) spikes and train of spikes/polyspikes in neocortex (black traces) and hippocampus (blue traces). (**D**) In the mutant, interictal spike frequency was significantly higher in hippocampus (blue box), compared to those generated in neocortex (orange box) or generalized in both regions (green box) (F=46.65, degrees of freedom (df)=27). (**E,F**) Power spectrum analysis displayed increased activity in mutant, as emphasized in the representative magnified segments. (**G,H**) Mutant hippocampus demonstrated significant higher activity in the gamma frequency range (F=3.932, df=27; neocortex: F=26.93, df=27). Data is represented as mean ± SEM scatter plots; differences were considered significant at p<0.05; ns, not significant. Scalebars: 0.2s,20µV (B,C); 1s,50µV (E,F). See also Figure 1 – figure supplement 1.

### Mutant CA1 has elevated, acutely regulatable network excitability *in vitro*

To assess the physiological aspects of Pik3ca-driven epilepsy at the tissue level, we performed hippocampal extracellular field recordings in acute forebrain slices. First, we conducted these recordings in P16-20 CA1 pyramidal layer, across a potassium concentration gradient in the recording buffer ([K^+^]: 3 mM to 12 mM; Figure 2A-C). This standard model shifts wild-type neuronal network from a silent state to tonic spiking and then to bursting, followed by a depolarization block at very high levels of [K^+^], thus mimicking a seizure-like response (Filatov et al. 2011). Compared to controls, mutant slices exhibited a narrow range of hypersynchronous bursting and an elevated mean baseline measured from the integrated trace, especially from 8mM to 12mM [K^+^] (Figure 2B,C; Figure 2 – figure supplement 2A). Mutant slices also showed significantly enhanced relative peak amplitudes (at 8-9 mM [K^+^]; Figure 2 – figure supplement 2B).

**Figure 2:**
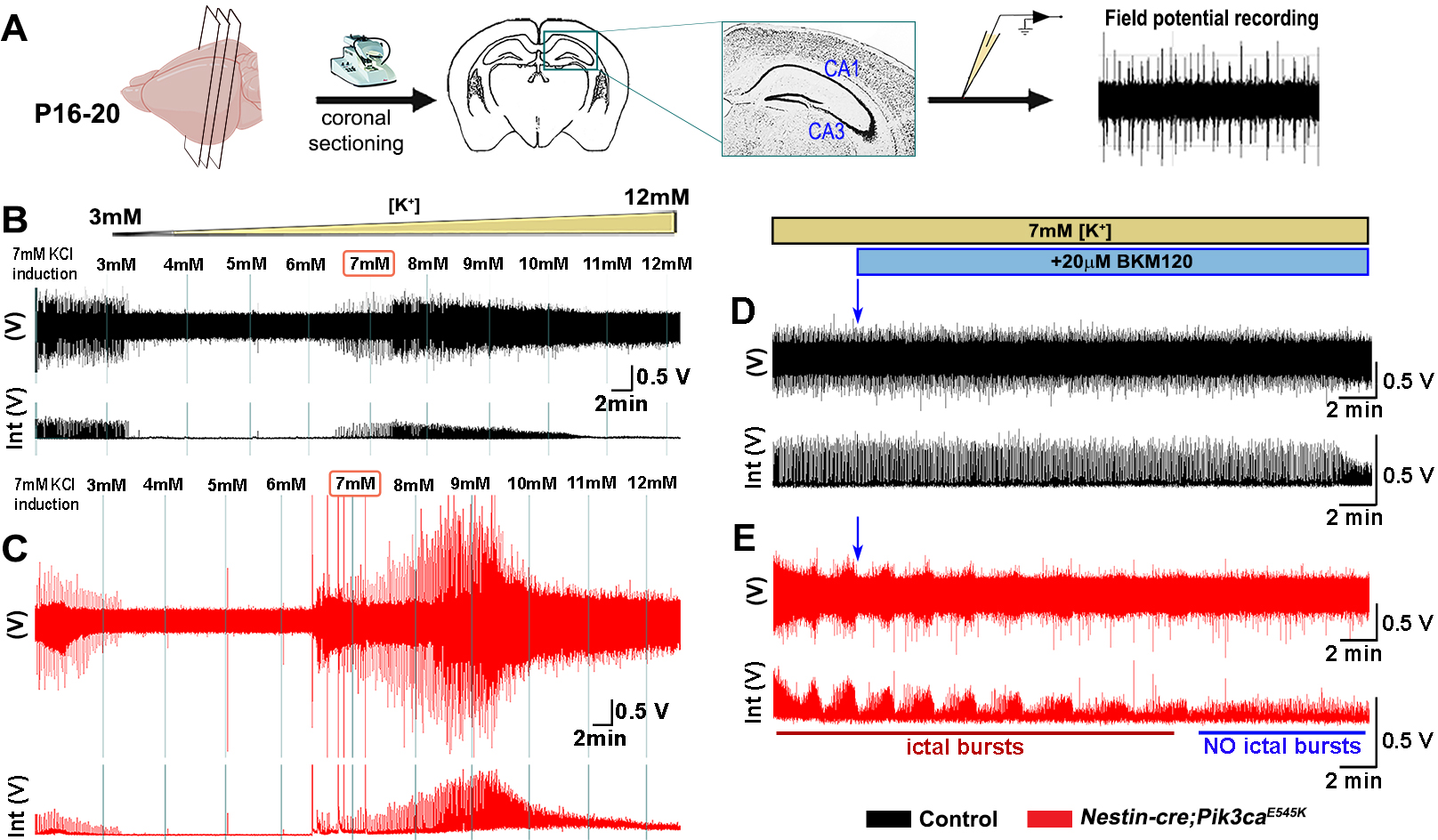
Extracellular mutant CA1 recordings demonstrate hyper-synchronization. (**A**) Schematized flowchart summarizes the acute coronal brain slicing for extracellular field potential recording. (**B,C**) Representative control (black) and mutant (red) voltage and integrated voltage traces across a [K^+^] gradient; comparison indicated significantly higher excitability in the mutant brain. (**D,E**) Acute extracellular BKM120 administration at constant 7mM [K^+^] aCSF suppressed synchronized ictal bursts in mutant CA1 field within ∼20-30min, with minimal effect on the control hippocampal slices. Scalebars: 2min,0.5V (B-E). See also Figure 2 – figure supplement 1.

Thus, the mutation lowers the excitability threshold and a depolarization block is reached faster than in controls. Further, acute extracellular application of BKM120 suppressed the hypersynchronous ictal bursts in the mutant CA1 field in 20-30 min, with minimal physiological effect on control brains (Figure 2D,E). These data provide evidence of a Pik3ca-driven active mechanism.

### Mutant CA1 and CA3 pyramidal neurons are hyperexcitable

Next, we investigated mechanisms driving the mutant epileptiform activity at the cellular level, using whole-cell patch-clamp recordings from CA1 and CA3 pyramidal neurons in P16-20 control and mutant brains (Figure 3A). Silent and spontaneously firing, tonic and burst-generating neurons were detected in both control and mutant slices (Figure 3B,C). However, mutant slices had a significantly higher proportion of burst-generating cells in both regions and lower proportion of tonic-firing cells in mutant CA1 compared to controls (Figure 3D,E). Additionally, we observed significantly higher tonic spike frequencies in mutant CA1 and CA3 and higher burst frequency in mutant CA1, compared to controls (Figure 3F,G). Burst-generating cells demonstrated multiple burst types. Specifically, we identified burst clusters and two types of plateau-bursts: paroxysmal depolarization shift (PDS) and non-PDS waveforms (Figure 3 – figure supplement 1A). We defined burst cluster as a multi-spike burst activity of random nature without a plateau potential; these were only observed in mutant slices (Figure 3 – figure supplement 1B). Plateau-bursts with depolarization shift resulting in sodium-spike inactivation were termed as PDS “bursting cells”. These depolarization shifts have previously been implicated as the intracellular correlate of *in vivo* interictal spikes (Marcuccilli et al. 2010; McCormick and Contreras 2001; Kubista, Boehm, and Hotka 2019; Tryba et al. 2019). We defined non-PDS bursts as those where plateau potential developed in absence of prominent sodium-spike inactivation. No significant differences between control and mutant hippocampal pyramidal cells were observed with respect to average burst duration or inter-burst interval (Figure 3 – figure supplement 1C-E).

**Figure 3:**
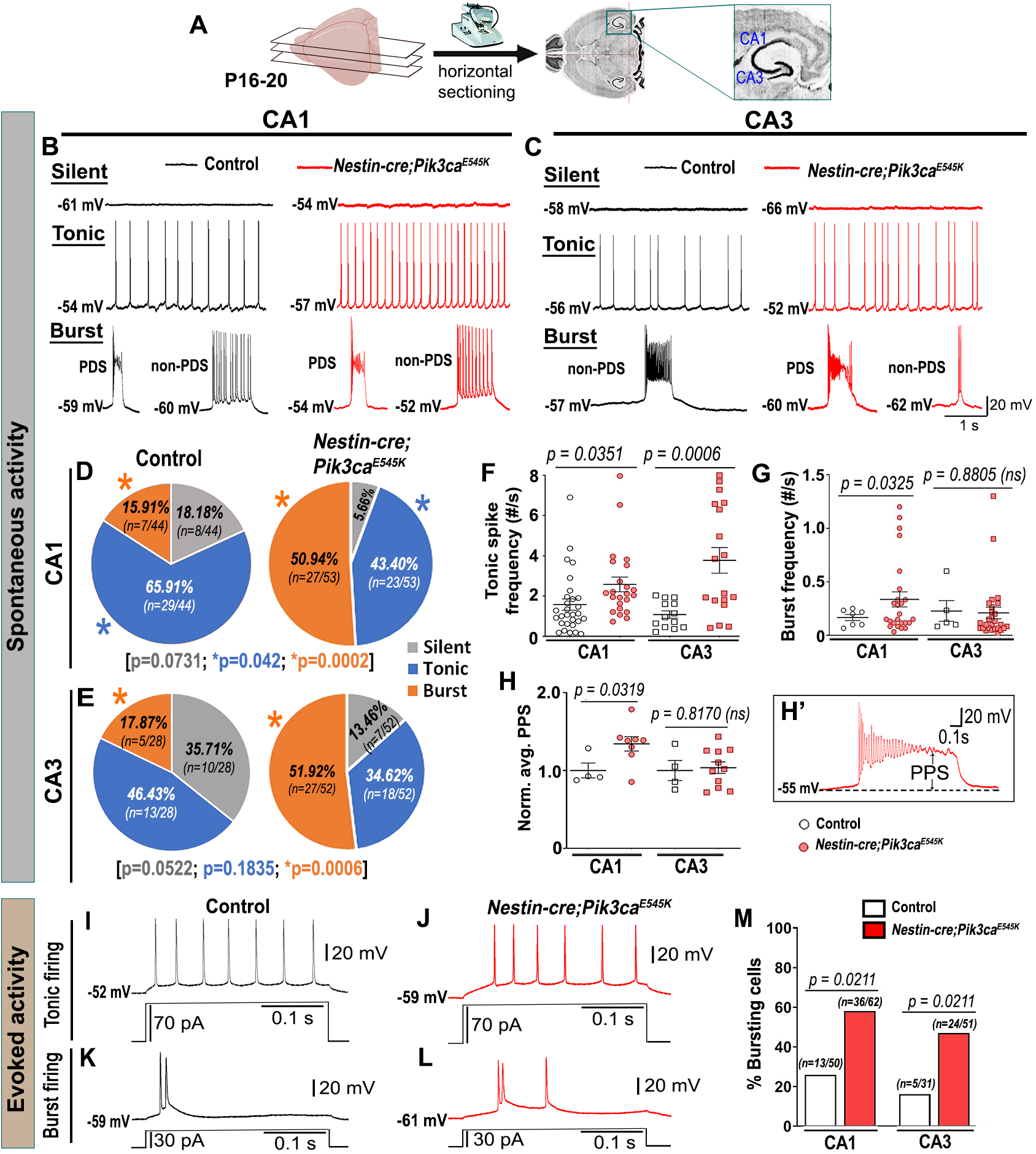
Mutant hippocampal neurons produce increased epileptiform burst activity. (**A**) Flowchart shows acute horizontal brain slicing for whole-cell recording. (**B-E**) Traces represent silent, tonic, and burst categories of CA1 and CA3 neurons based on spontaneous cellular activity; respective pie charts marked proportion of recorded cells. Mutant CA1 and CA3 exhibited significantly higher proportions of burst-firing cells compared to controls. Significantly fewer tonic-firing cells were observed in mutant CA1. (**F,G**) relative to controls, spontaneous tonic spike frequencies were significantly higher in mutant CA1 (t=2.175, df=43.71) and CA3 (t=4.063, df=19.78) cells; burst frequency was significantly higher in mutant CA1 (CA1: t=2.244, df=29.38; CA3: t=0.1561, df=6.754). (**H**) Plateau potential shift (PPS) in mutant bursting cells, as depicted in (**H’**), was significantly higher in CA1 (t=2.586, df=8.118) but similar in CA3 (t=0.2436, df=5.110), compared to respective controls. (**I-L**) Representative evoked voltage traces of control and mutant showed tonic and burst firing in response to current steps. (**M**) Evoked recording also marked significantly higher bursting cell proportions in the mutant. Data is represented as pie charts, % bar graphs and mean ± SEM scatter plots; differences were considered significant at p<0.05; ns, not significant. Scalebars: 1s,20mV (B,D); 0.1s,20mV (H’, I-L). See also Figure 3 – figure supplements 1-2.

However, the plateau potential shift (PPS), defined here as the difference of the steady state plateau potential and the resting membrane potential (RMP), was significantly larger in mutant CA1 but not in CA3, relative to respective controls (Figure 3H,H’). This was despite similar RMP across control and mutant hippocampal neurons (Figure 3 – figure supplement 2A-C). The evoked current-clamp recordings further validated the spontaneous activity results, especially by displaying significantly higher percentage of burst-generating pyramidal neurons in the mutant hippocampus (Figure 3I-M).

Incidentally, no significant difference in membrane intrinsic properties like input resistance, rheobase current and burst-threshold current, as well as evoked tonic spike frequencies (for the tested 0-90pA range), was observed between control and mutant cells (Table 1; Figure 3 – figure supplement 2D-F). But the decay time constant for mutant CA3 neurons was significantly longer than that in mutant CA1 and control groups (Table 1). With similar resistance, this implied that mutant CA3 neurons have higher membrane capacitance than other cell groups. Together, these data demonstrate that Pik3ca overactivation results in intracellular hyperexcitability, with some distinct cell type-specific effects.

**Table 1:**
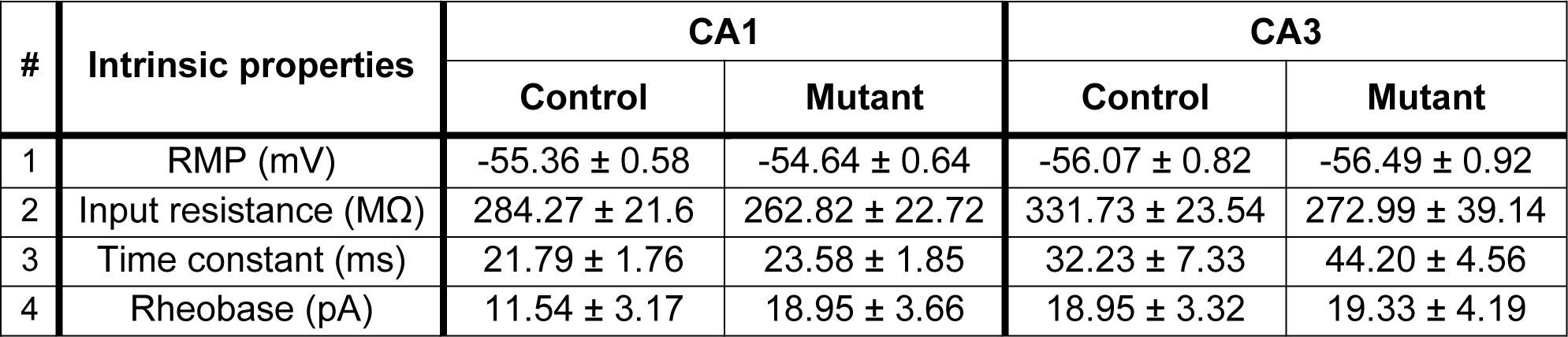
Summary table of intrinsic membrane properties. Summary of basic intrinsic membrane properties quantitated and compared between control and mutant CA1 and CA3 neurons. No significant differences in RMP, input resistance and rheobase current (minimum injected current required to elicit the first action potential) were observed between control and mutant neurons, in both CA1 and CA3. Decay time constant of mutant CA3 neurons was significantly longer than that of control CA1 (p < 0.0001) and CA3 (p = 0.0071) cells, as well as of mutant CA1 neurons (p < 0.0001) (F=31.21, df=33). Data is represented as mean ± SEM; differences were considered significant at p<0.05.

To determine whether the Pik3ca-related hyperexcitability is driven by altered synaptic interactions or other intrinsic properties, we assessed the effects of channel and receptor blockers on mutant hippocampal physiology. Blocking glutamatergic inputs by extracellular administration of NMDA and non-NMDA receptor-antagonists, 3-((±)2-carboxypiperazin-4yl) propyl-1-phosphate (CPP) and 6-cyano-7-nitroquinoxaline-2,3-dione (CNQX) respectively, had no overt physiological effect on the majority of mutant hippocampal neurons (Figure 4 – figure supplement 1A-C,E,F,H-K). Similarly, blocking inhibitory synaptic inputs with gabazine did not significantly alter the mutant firing patterns, spike frequencies or PPS (Figure 4 – figure supplement 1A,D,G,L). The proportion of mutant CA1 neurons affected by these channel blockers was comparatively less than that in CA3. Our data lead to the important conclusion that Pik3ca-driven epileptiform activity is primarily not dependent on synaptic transmission. In contrast, inhibition of calcium (Ca^2+^)-dependent inward current by extracellular cadmium (Cd^2+^) attenuated the paroxysmal bursts and reduced burst frequency and PPS (Figure 4A-D), indicating a calcium channel-dependent mechanism underlying the Pik3ca-dependent epileptiform activity. Intracellular cesium blocked potassium channels and related currents, altering the intrinsic firing pattern in both control and mutant hippocampal slices. Specifically, compared to regular baseline recordings, intracellular cesium considerably reduced the proportion of tonic-firing cells in both CA1 and CA3 (Figure 4E-H, compare to Figure 3D,E). Unlike the spontaneous recordings, intracellular cesium prompted the burst frequency in mutant CA1 to normalize and in mutant CA3 to significantly rise, relative to respective controls (Figure 4I, compare to Figure 3G).

**Figure 4:**
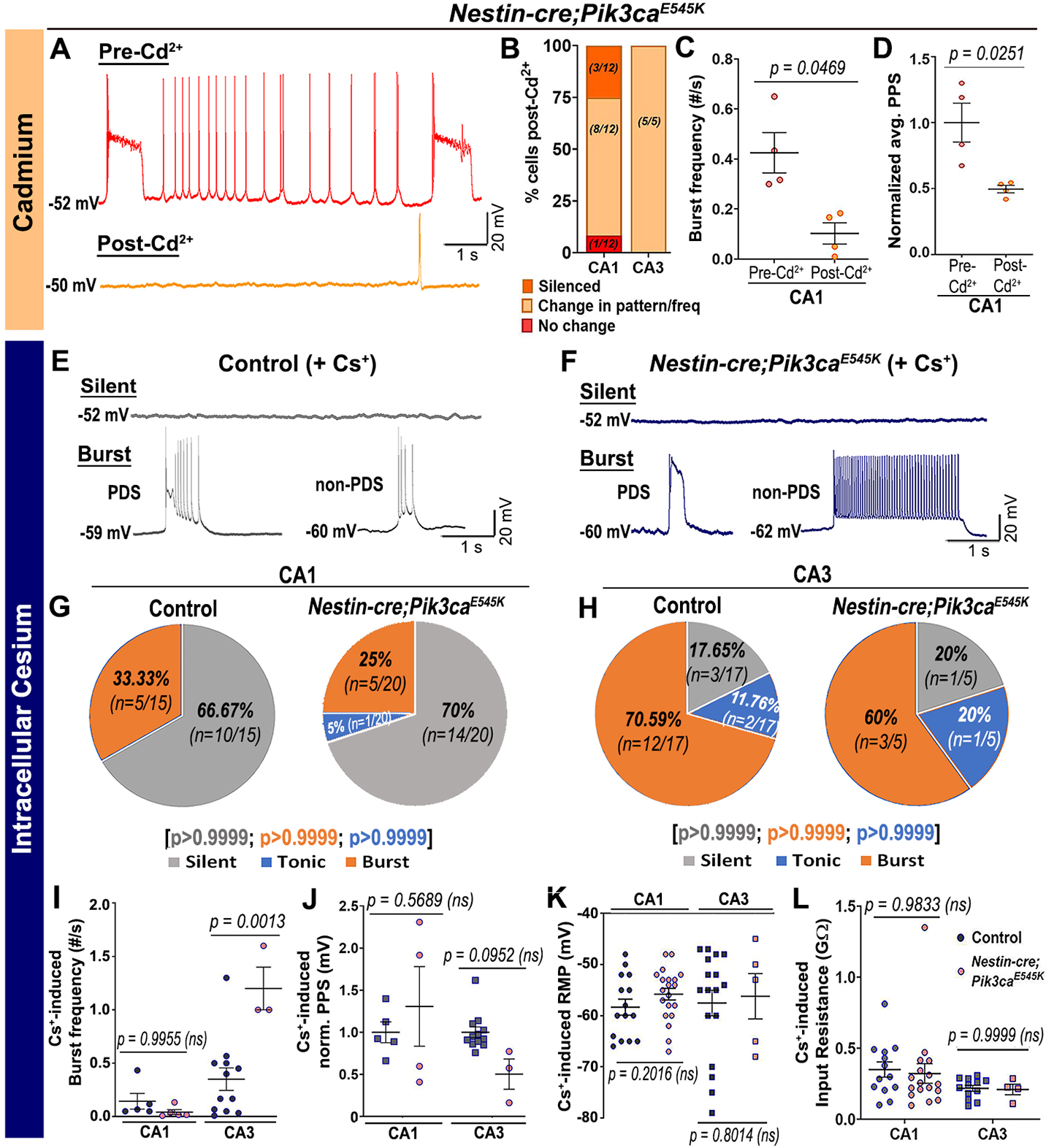
Pik3ca-related burst characteristics are dependent on cell-intrinsic calcium and potassium channel regulation. (**A-D**) Blocking calcium channels by extracellular cadmium (Cd^2+^) significantly altered intrinsic bursting pattern of mutant neurons including burst frequency (t=3.267, df=3) and PPS (t=4.172, df=3). (**E-H**) Representative traces and respective pie charts show types of cellular activity in control and mutant CA1 and CA3 neurons after administering cesium intracellularly. In response to cesium, tonic firing cells were significantly reduced and original differences in CA1 and CA3 cell proportions between control and mutant were normalized. (**I**) Cs^+^-induced burst frequency was significantly higher in mutant CA3 than those in respective controls (F= 26.31, df=21); unlike normal spontaneous state, no such difference was observed in CA1. (**J-L**) Intracellular cesium lowered the spontaneously enhanced mutant PPS in CA1 to control levels (F=4.003, df=20), while having no effect on RMP (CA1: t=1.308, df=27.32; CA3: t=0.2616, df=6.816) or input resistance (F= 2.936, df=42) in both control and mutant neurons. Data is represented as pie charts, % bar graphs and mean ± SEM scatter plots; differences were considered significant at p<0.05; ns, not significant. Scalebars: 1s,20mV (A,E,F). See also Figure 4 – figure supplement 1.

Intracellular cesium also normalized the mutant PPS to control levels (Figure 4J, compare to Figure 3H). No overt effect on RMP or input resistance was seen in cesium-treated control and mutant cells (Figure 4K,L). We conclude that Pik3ca-related epileptiform activity is primarily caused by a complex set of altered non-synaptic cell-intrinsic properties.

### Acute treatment of epileptiform activity with PI3K pathway drugs

The acute suppression of epileptiform activity in *Nestin-cre;Pik3ca^E545K^* mice by BKM120, both *in vivo* (Roy et al. 2015) and at the tissue level, prompted us to dissect the downstream pathway dynamics using inhibitors at the cellular level, in order to coarsely determine their mechanistic roles and identify new therapeutic targets to treat intractable epilepsy (Figure 5A). Acute extracellular administration of BKM120 or the pan-AKT inhibitor AZD5363 (Davies et al. 2012) resulted in a large-scale and significant alteration of firing pattern, frequencies and a relative reduction of PPS, sometimes leading to gradual silencing of the recorded neurons (Figure 5B-I,I’). Neither drug had any significant effect on neuronal RMP or tonic rheobase current (Figure 5 – figure supplement 1A,B,E,F). In contrast to BKM120, acute AKT downregulation by AZD5363 significantly reduced the mutant burst duration and inter-burst interval (Figure 5 – figure supplement 1C,D,G,H). Further, AZD5363 altered the physiology of all mutant CA3 neurons much faster (∼8 min) than BKM120 (∼32 min); however, both drugs changed the intracellular activity of ∼50% cells within 4 min (Figure 5 – figure supplement 1I,J).

**Figure 5:**
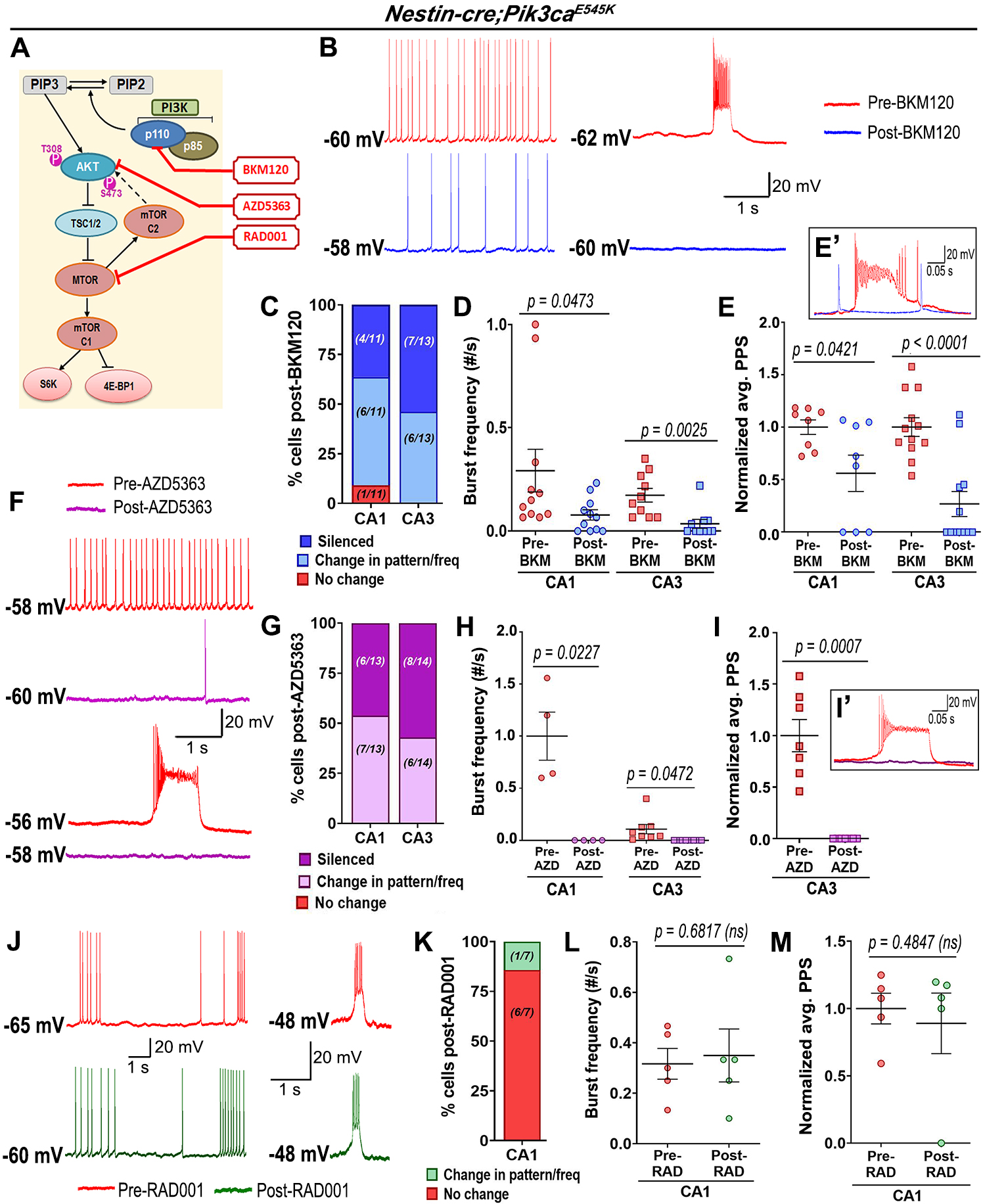
Acute inhibition of Pik3ca or Akt suppresses hyperactivity in mutant cells. (**A**) Schematic of simplified PI3K-AKT-MTOR pathway, marking targets of action for BKM120, AZD5363 and RAD001. Representative traces demonstrate differential action of BKM120 (**B**), AZD5363 (**F**) and RAD001 (**J**) on *Nestin-cre;Pik3ca^E545K^* hippocampal neurons, ranging from being unaffected to change in frequency/pattern to silencing, as quantitated respectively in (**C,G,K**). (**D,E,H,I**) BKM120 and AZD5363 significantly reduced mutant burst frequency (BKM120: F=8.937, df=38; AZD5363: F=43.50, df=20) and PPS (BKM120: F=24.39, df=36; AZD5363: t=6.386, df=6) in both CA1 and CA3. (**E’,I’**) Insets show activity traces of mutant bursting cell before (red) and after BKM120 (blue) or AZD5363 (pink) administration. (**L,M**) Acute RAD001 treatment had no overt effect on the mutant burst frequency (t=0.4414, df=4) or PPS (t=0.7692, df=4). Data is represented as % bar graphs and mean ± SEM scatter plots; differences were considered significant at p<0.05; ns, not significant. Scalebars: 1s,20mV (B,F,J); 0.05s,20mV (E’,I’). See also Figure 5 – figure supplements 1-2.

AZD5363 also blocked spontaneous bursts in mutant neurons faster than BKM120 (Figure 5 – figure supplement 1K). Thus, acute regulation of both PI3K and AKT activity directly suppressed *Pik3ca^E545K^*-driven epileptiform neuronal activity. In contrast, acute administration of the MTOR inhibitor RAD001 (everolimus) (Cho 2011; Krueger et al. 2010) showed no overt effect on the mutant RMP, tonic or burst firing patterns, frequencies, average PPS, burst duration, inter-burst interval and rheobase current (Figure 5J-M, Figure 5 – figure supplement 2A-D). This was despite western blot confirmation that acute treatment of forebrain slices with RAD001 modulated direct downstream targets of MTOR (Figure 5 – figure supplement 2E-G). In summary, our study demonstrates that epileptiform activity caused by Pik3ca overactivation is based on multiple intrinsic electrophysiological characteristics that are acutely modifiable by downregulating PI3K and AKT activity, but not MTOR.

## DISCUSSION

Activating mutations in the PI3K-AKT-MTOR pathway commonly cause pediatric intractable epilepsy (Kim and Lee 2019; Dobyns and Mirzaa 2019); yet the underlying mechanisms remain largely undefined. Using our *Nestin-cre;Pik3ca^E545K^* model, we now establish that Pik3ca-dependent neuronal hyperexcitability in hippocampal pyramidal neurons is driven primarily by intrinsic neuronal mechanisms. Acute attenuation of epileptiform activity by inhibiting AKT but not MTOR also reveals a previously unknown role of PI3K signaling in neuronal homeostasis. Our mouse model of human intractable epilepsy is now validated across whole animal, tissue, and cellular levels.

Epileptiform activity is typically marked by increased tonic depolarizations and/or quasiperiodic bursts, mediated by intrinsic membrane properties, ephaptic or synaptic/circuit-level interactions (McCormick and Contreras 2001; Dudek, Yasumura, and Rash 1998). We confirmed enhanced interictal spike frequency and diverse, synchronized epileptiform spike patterns in the mutant brains *in vivo*, predominantly in the hippocampus. This is consistent with studies of *PIK3CA* variants in brain tumors and mouse models with patient-related *Pik3r2* mutations, which are less common causes of intractable epilepsy (Yu et al. 2020; Shi et al. 2020). Our *in vitro* analyses revealed intracellular correlates of epileptiform activity and enhanced firing frequencies in the mutant hippocampal pyramidal neurons. Relative to controls, *Pik3ca^E545K^* neurons exhibited significantly higher proportions of bursting cells and higher heterogeneity in burst patterns, including burst clusters and PDSs. A prior study showed that human neurons from surgically resected neocortical samples of pediatric patients with focal cortical dysplasia (FCD) exhibited remarkably similar burst characteristics (Marcuccilli et al. 2010). These samples were not genotyped; however, since most cases of FCD and epilepsy result from variants in PI3K pathway genes (Mirzaa 2018), such cross-species phenotypic correlation is remarkable and emphasizes the clinical relevance of our mouse model.

The PI3K-AKT-MTOR pathway is also known to influence synaptogenesis and neuronal plasticity (Sanchez-Alegria et al. 2018). For example, neocortical samples from patient and mouse models with *TSC* mutations exhibit synaptic input-mediated epileptogenic hyperexcitability (Wang et al. 2007). However, blocking glutamatergic (NMDA and non-NMDA) or GABAergic synaptic inputs showed no significant effect on the neuronal activity in our mutant brain slices. We acknowledge that altered neuronal circuitry may be relevant to Pik3ca-mediated epilepsy, but our data demonstrate that altered synaptic interactions are at least not primary mediators of dynamic hippocampal neuronal hyperexcitability.

Significant reduction of burst-generating cells, burst frequency and PPS by blocking voltage-dependent calcium or potassium channels indicate that *Pik3ca^E545K^*-driven epileptiform activity is predominantly mediated by multiple altered cell-intrinsic properties. Yet, some cell-intrinsic properties, such as burst frequencies, PPS and time constants, are distinctly different between mutant CA1 and CA3 neurons. Since all neurons do not react uniformly to either pathway overactivation or Cd^2+^/Cs^+^-dependent channel inhibition, current anti-seizure drugs that typically target single ion channels cannot address the heterogeneous dysregulation, thus potentially explaining epileptic intractability in this patient population. It is intriguing to note that intercellular signaling heterogeneity involving PI3K-AKT-MTOR pathway was also reported by a recent quantitative study that highlighted its potential relevance to human disorders, such as insulin resistance (Norris et al. 2021).

PI3K-AKT-MTOR inhibitors developed to treat cancer potentially represent novel anti-seizure therapeutics (Cardamone et al. 2014; Lindhurst et al. 2015; Maira et al. 2012; Zou et al. 2020; Forde et al. 2021; Venot et al. 2018). Rapamycin and its analogs, that are currently in clinical use to treat epilepsy in TSC patients, curbed epileptiform activity in mouse models harboring *TSC1/2*, *RHEB, MTOR* or *PTEN* mutations when administered long-term (>7-day) (Curatolo and Moavero 2013; Crino 2016; Lasarge and Danzer 2014; Stafstrom 2019; Kim and Lee 2019). In contrast, we show acute inhibition of epileptiform activity by BKM120 *in vitro*, confirming our previous *in vivo* finding (Roy et al. 2015). We also demonstrate that acute AKT inhibition with AZD5363 has similar effects. This first preclinical study using AZD5363 to treat epilepsy shows that acute Pik3ca-driven epileptiform mechanism is predominantly AKT-dependent, limiting roles for other PI3K targets. Intriguingly, rapamycin analog RAD001 had minimal acute effect in our model. While elevated MTOR activity is known to be epileptogenic (Crino 2016; Kim and Lee 2019), our data suggest that alternate PI3K-AKT targets acutely regulate neuronal hyperexcitability. This study establishes an important foundation to determine active pathway-driven cellular mechanisms and to assess the plethora of available PI3K pathway inhibitors, facilitating new molecularly rational therapeutic interventions for intractable epilepsy.

## MATERIALS AND METHODS

### Mice

The following mouse lines were used: *Nestin-cre* (Jackson Labs, Bar Harbor, Maine, USA; Stock #003771, RRID: IMSR_JAX:003771), conditional *Pik3ca^E545K^* knock-in (Robinson et al. 2012). *Nestin-cre* and *Pik3ca^E545K^* mouse lines were maintained in C57BL/6 and FVB backgrounds respectively. We have designated *Nestin-cre;Pik3ca^E545K^* conditional mutant mice as “*Pik3ca* mutants” or “mutants” throughout the manuscript.

All mice were housed in Optimice cages with aspen bedding at the Seattle Children’s Research Institute’s specific pathogen-free (SPF) vivarium facility (light “ON”: 6am – 8pm). Noon of the day of vaginal plug was designated as embryonic day 0.5 (E0.5). The day of birth was designated as postnatal day 0 (P0). Genotyping by PCR was done using separate sets of primers for the *Cre* coding region and the *Pik3ca^E545K^* allele, as previously described(Roy et al. 2015). All mouse procedures were approved and conducted in accordance with the guidelines laid down by the Institutional Animal Care and Use Committees (IACUC) of Seattle Children’s Research Institute, Seattle, WA, USA. ARRIVE guidelines have been followed for reporting work involving animal research.

### *In vivo* electrophysiology

5 control and 6 *Nestin-cre;Pik3ca^E545K^* mutant mice (age: ∼P70) were used for *in vivo* regular and depth-electrode electrophysiology experiments. We saw no sex-dependent data correlation (Roy et al. 2015).

*Electrode implantation surgery:* Mice underwent survival surgery to implant ECoG, EMG, and hippocampal depth electrodes under isoflurane anesthesia using similar procedures as reported in our prior work (Bolea et al. 2019; Roy et al. 2015). The ECoG electrodes consisted of silver wire (diameter: 130 µm bare; 180 µm coated) attached to micro screws. The ECoG electrodes were implanted bilaterally through small cranial burr holes above the somatosensory cortices. A similar reference electrode was placed above the cerebellum following the same procedure. The depth electrodes were made of 2 fine twisted tungsten wires (30um nylon coated, California fine wire) and implanted in the hippocampal CA1 region (coordinates: -2.0 mm anteroposterior, 1.5 mm mediolateral, 1.9 mm dorsoventral in reference to bregma) to record local field potential (LFP). All electrodes were connected to interface connector and fixed to the skull with dental cement (Lang Dental Manufacturing Co., Inc., Wheeling, IL). Mice were allowed to recover from surgery for 2-3 days.

*Video, electroencephalagraphy and local field potential (Video-EEG-LFP) recording:* Simultaneous video-EEG-LFP recordings were collected from conscious mice on PowerLab 8/35 and 16/30 data acquisition units, using LabChart 7.3.3 software (AD Instruments, Colorado Spring, USA). ∼6 hours of baseline ECoG tracings were visually reviewed for the presence of spontaneous epileptiform events, as previously studied (Kalume 2013; Liautard et al. 2013; Roy et al. 2015). All bioelectrical signals were acquired at a 20 KHz sampling rate. The ECoG signals were processed with a 1-70 Hz bandpass filter and the LFP signal with a 5 Hz high-pass filter. Interictal spikes were identified as transient, clearly distinguished from background activity, with pointed peak and slow wave. Myoclonic seizures were identified as shock-like muscular jerks on video, associated with a spike or polyspike-wave complex on EEG. Power spectral analysis and visual inspection of the data were conducted to characterize the EEG activity in different frequency bands and identify epileptiform event on the ECoG and local CA1 field recordings using the Labchart 8.2 software.

### *In vitro* electrophysiology

*Slice preparation:* Postnatal (P16-20) pups were anesthetized briefly in a closed chamber by administering isoflurane (5% flow rate) or CO2 (constant flow rate: 10-30% of chamber vol/min); then perfused transcardially with ice-cold low Na^+^-buffer (“slicing solution,” which included the following: 252 mM sucrose, 2 mM KCl, 2 mM MgCl2, 2.6 mM CaCl2, 1.2 mM NaH2PO4, 26 mM NaHCO3, and 15 mM glucose, with the pH adjusted to 7.4 and the osmolarity to 310 ± 5 mOsm). Brain was dissected out by separating the head and cutting along the skull sutures using fine scissors and forceps. The forebrain was isolated in ice-cold, oxygenated (95% O2, 5% CO2) slicing solution. A slanted (∼15° from vertical) agar block was secured on a specimen tray as a support for the brain during slicing. The isolated forebrain was glued with cyanoacrylate in the orientation depending on the type of slices we wanted; and then placed in the vibratome for slicing. Slicing was proceeded until hippocampal landmark was visible. For coronal slices (extracellular recordings), the forebrain was placed such that its antero-posterior (A-P) axis was perpendicular to the specimen tray/vibratome blade, and olfactory bulb distal to the tray surface. For horizontal slices (whole-cell recordings), the forebrain was oriented such that it’s A-P axis was parallel to the specimen tray and ventral side (hippocampal side) was on the top, distal to the tray surface. Once the hippocampus became visible, thick, acute slices (thickness: 350μm for extracellular recordings, 200-250μm for whole-cell patch clamp recordings) were taken and incubated in ice-cold slicing solution continuously oxygenated with 95% O2/5% CO2. After cutting, slices were immediately returned to the same solution and maintained in a warm bath (28 ± 0.5°C) for recovery. After 30 min, they were transferred into regular artificial cerebrospinal fluid (aCSF), composed of the same components as slicing solution except for the replacement of sucrose with 126 mM NaCl. Slices were then kept at room temperature, continuously superfused with oxygenated aCSF, until recording. Given that *Pik3ca^E545K^* mutant is megalencephalic, a greater number of hippocampal/forebrain sections were obtained. However, care was taken so that the dorso-ventral plane(s) used for recording was always comparable between control and mutant slices. No randomization was used. Tissue collection was not performed blind since the mice were subjected to genotyping and drug administration.

*Recording:* Slices were transferred into a recording chamber, continuously superfused with oxygenated aCSF, for extracellular or intracellular whole-cell recordings. The pClamp software suite (Molecular Devices; RRID:SCR_011323) was used for data acquisition and analysis. Signals were amplified (MultiClamp 700A, Axon Instruments, Molecular Devices, USA), digitized (D1322A, Axon Instruments, Molecular Devices, USA), and stored in a computer for post-hoc analysis.

*a) Extracellular recording:* To determine if hippocampal neuronal populations in *Pik3ca^E545K^* mutants have different electrochemical signatures than controls, extracellular population recordings were performed on CA1 hippocampal pyramidal neuronal layer in acute 350 µm-thick coronal brain slices of ∼P16-20 *Pik3ca^E545K^* mutant and control littermates (n=6). To induce epileptiform activity in the slices, we increased potassium concentration ([K^+^]) (Borck and Jefferys 1999; Leschinger et al. 1993) from 3 mM to 12 mM, in 1mM increments at 30°C. This standard model induces wild-type neurons to transition from a silent neuronal state to tonic spiking and then to bursting, followed by depolarization block at very high [K^+^] (Filatov et al. 2011).
*b) Intracellular recording:* Horizontal hemi-forebrain slices with prominent hippocampus were used for intracellular whole-cell visual patch-clamp experiments. Slices transferred to the recording chamber were maintained at 30-34°C, constantly superfused with oxygenated aCSF. Borosilicate glass capillaries/pipettes for patch-clamp recording had electrode resistance (Re) optimally kept around 5-8 MΩ, after being filled with internal solution containing the following: potassium gluconate (∼132 mM), KCl (5 mM), HEPES (10 mM), EGTA (5 mM), CaCl2.H2O (0.5 mM), MgCl2 (2 mM), disodium phosphocreatine (5 mM), disodium-ATP (4 mM), trisodium-GTP (0.5 mM), EGTA (5 mM). Cells were visualized under brightfield optics using the 40X water-immersion objective of an upright microscope (Olympus, BX51WI). The patch electrode was advanced toward the target cells by a micromanipulator (MP-225, Sutter Instrument Company, USA) and 1 GΩ seal was established, typically by a small negative pressure, with the membrane ruptured by gentle suction and/or zap pulses. Whole-cell patch-clamp recording was performed from cell bodies of pyramidal neurons of hippocampal CA1 and CA3 respectively. Following intracellular recording protocols were also used:
i. Spontaneous gap-free recordings in current (I)-clamp: cells were typically tested at resting conditions (without current injection) unless noticed otherwise.
ii. Evoked I-clamp steps protocol: current steps start at -50 pA, incremental 10 pA, duration; 300 ms; 15 steps (-90 to +50 pA) were recorded across experiments.

### Acute chemical assays

#### Channel blockers

Chemicals inhibiting specific ion channels were introduced in the bath/recording chamber during *in vitro* whole-cell recordings to compare different components contributing to the neural activity in control and *Pik3ca^E545K^* mutant mice. To block all fast-synaptic excitatory transmission, 3-((±)2-carboxypiperazin-4yl) propyl-1-phosphate (CPP, NMDA receptor antagonist, 20 μM; Tocris Bioscience, UK) and 6-cyano-7-nitroquinoxaline-2,3-dione (CNQX, AMPA/kainite (non-NMDA) receptor antagonist, 20 μM, diluted in DMSO; Alomone Labs, Israel) were introduced in the bath (Lehmann et al. 1987; Lester et al. 1989). SR95531 hydrobromide (Gabazine, GABAA receptor antagonist, 10 μM, Tocris Bioscience, UK) were used to block inhibitory synaptic transmission. Divalent cation Cd^2+^ (CdCl2, 100 μM; Sigma-Aldrich, USA) (Boissard et al. 2002) was used to depress synaptic transmission and block inward calcium-selective current, isolating the outward K^+^ current. To block voltage-dependent K^+^ channels, intracellular administration of cesium (Covey, Kauer, and Casseday 1996) was done by replacing potassium gluconate with cesium gluconate in the internal solution while maintaining the same osmolarity.

#### PI3K pathway drugs

Drugs regulating PI3K pathway were added to the circulating recording buffer, thus getting acutely administered in the brain slice placed in the recording chamber. For protein analyses, these were added at identical concentration to the brain slices incubated in oxygenated aCSF for ∼1hr. The following PI3K pathway inhibitors were dissolved in 100% DMSO and used *in vitro*: pan-PI3K inhibitor BKM120 (Buparlisib; Novartis, Switzerland; working concentration: ∼3.5 µM); pan-AKT inhibitor AZD5363 (Capivasertib; Selleckchem, USA; working concentration: 0.5 µM); MTOR inhibitor RAD001 (Everolimus, Chem Express Cat# 159351-69-6; working concentration: 0.52 µM).

### Western blotting

Horizontal forebrain slices from 5 control and 5 mutant mice were obtained and recovered in oxygenated aCSF as detailed above. Slices showing nice hippocampal morphology were selected, treated for 1 hour with DMSO, BKM120, AZD5363 and RAD001 respectively, in the same concentrations used for recording. The treated slices were then flash-frozen in liquid nitrogen and stored at -80°C for post-processing. These brain slices were lysed in NP-40 lysis buffer (150 mM NaCl, 50 mM Tris (pH 7.4), 1% NP-40, 10 mM NaF, 2 mM sodium orthovanadate + protease/phosphatase inhibitor cocktails [Sigma, USA]). Samples were normalized to equal protein concentrations (0.333 mg/ml) using Pierce BCA protein assay (Thermo Fisher Scientific, USA).

Standards were made through serial dilutions of 10 mg/ml BSA. Samples were diluted into a final concentration of 1x Laemmli sample buffer and boiled at 95°C for ten minutes. 7.5% 10-well gels were prepared, and samples were run at 120V for 1.5 hours. Gels were transferred onto polyvinylidene difluoride (Millipore) membranes and run either overnight at 25V or for 2 hours at 60V. Membranes were blocked in 4% milk in TBST (0.05 M Tris, 0.15 M NaCl, pH7.2, 0.1% (v/v) Tween20) for one hour at room temperature, and primary antibodies were applied overnight at 4°C in blocking medium.

Primary antibodies for western blots were diluted as follows: rabbit anti-Phospho-AKT Ser473 (D9E) (Cell Signaling Technology, USA; RRID: AB_2797780; 1:2000), mouse anti-Pan-AKT (40D4) (Cell Signaling Technology, USA; Cat# 2920; 1:2000), rabbit anti-Phospho-S6 Ribosomal Protein Ser235/236 (Cell Signaling Technology, USA; RRID: AB_2721245, 1:2000), mouse anti-S6 Ribosomal Protein (54D2) (Cell Signaling Technology, USA; RRID: AB_2238583, 1:1000), rabbit anti-beta(b)-Actin (GeneTex, USA; RRID: AB_1949572, 1:10000). After washing and probing respectively with goat anti-rabbit (RRID: AB_2313567) and anti-mouse (RRID: AB_10015289) horseradish peroxidase (HRP)-conjugated secondary antibodies (1:10000; Jackson ImmunoResearch Labs, USA), blots were imaged using Femto chemiluminescent detection reagents (Thermo Fisher Scientific, USA; Cat# 34095) in a FluorChem R western blot imaging system (ProteinSimple, Bio-Techne, USA). 8-bit images were used as a representative western blot. 16-bit images were used to quantify the intensity of each band using ImageJ v1.53. Regions of interest were drawn in each sample’s lane. After quantifying the lane in a histogram, the peak representing the band of interest was isolated and the area of the region was measured as the band’s quantification.

### Quantitative and Statistical Analyses

Number of mice used was consistent with previous experiments completed and published by us and other investigators. For extracellular field potential analyses, baseline potential from integrated traces were measured for each [K^+^] and normalized to the value at 3 mM for each slice per genotype; relative peak amplitude was similarly measured for control and mutant slices for 3,7,8,9 mM of [K^+^]. In whole-cell patch-clamp recording, plateau potential shift (PPS) was calculated by subtracting the baseline potential from the highest plateau/burst potential in a paroxysmal depolarizing event. Burst duration and inter-burst interval were measured as shown in Figure 3 – figure supplement 1 and averaged across multiple bursts per cell. Input resistance was measured from evoked current-clamp recordings by dividing the voltage difference (measured at -10pA to -30pA I-steps) by the current interval (20pA). The decay membrane time constant was obtained by recording the membrane response to 20pA hyperpolarizing current pulses (300 ms duration, 1 Hz) and fitting the response to a single exponential curve. We chose 20pA of hyperpolarizing current because such a current intensity did not produce sag. Evoked spike frequencies and rheobase current were calculated exclusively from tonic-firing cells. Burst threshold current was calculated from burst-generating cells as the first current step which induced burst. *In vitro* electrophysiological analyses were performed using Clampfit 10.7 and 11 (pClamp, Molecular Devices, USA). For Western Blots, each band intensity was recalculated relative to its respective b-actin band intensity, and then normalized across average intensity per protein lane. Normalized ratios of phosphoproteins over total proteins were thereafter obtained.

Statistical significance was assessed using 2-tailed unpaired t-tests with Welch’s correction (EEG power analyses, whole-cell RMP, burst duration, inter-burst interval, Cd^2+^ data), 2-tailed paired t-tests (field potential baseline measurement, rheobase current, drug-treated analyses) and ANOVA followed by Tukey post-tests (EEG interictal spike frequency, field potential relative peak amplitude; whole-cell tonic spike and burst frequencies, cell proportions, PPS, evoked spike frequency and time constant; Western blots). Normal distribution was assumed for the data analyses, when required. For EEG-ECoG-LFP experiments, quantitative and statistical data analysis was performed in Labchart 8.2 software (AD Instruments, Colorado Spring, USA) and Igor Pro 6.37 (WaveMetrics Inc., USA); final graphs were made in GraphPad Prism v7.0 (GraphPad Software Inc., San Diego, USA). For the remaining data, statistical analyses and graph plotting were done using GraphPad Prism v7.0 and Microsoft Excel.

Differences were considered significant at p<0.05.

## Acknowledgments

We thank Suzanne J. Baker for gifts of mouse lines (*Nestin-cre* and *Pik3ca^E545K^*); Leon Murphy (Novartis) for BKM120; Rory M. Murphy, Amanda P. Tran Hartman, Nikhil Sahai and Jiyun Ryu for technical assistance; Aguan D. Wei and Paul Wakenight for discussions. This work is funded by the NIH grants 1R01NS099027 (K.J.M.), R01MH113545 (S.E.P.S.), R01NS102796 (F.K.), and CURE Sleep and Epilepsy Grant (F.K.).

## Author Contributions

A.R., F.K., J.M.R. and K.J.M. contributed to the study conception and design. A.R., V.H., A.M.B., D.T.W. and F.K. contributed to the data collection. A.R. and F.K. contributed to in vitro and in vivo data analyses respectively.

D.T.W. and S.E.P.S. contributed to Western blot data analyses and interpretation. A.R., F.K., J.M.R., K.J.M. contributed to the overall data interpretation. S.E.P.S., F.K. and K.J.M. provided the funding resources. A.R. wrote the first draft of the manuscript and others commented on previous versions of the manuscript. All authors read and approved the final manuscript.

## Ethics

Animal experimentation: All animal experimentation was conducted in accordance with the guidelines laid down by the Institutional Animal Care and Use Committees (IACUC) of Seattle Children’s Research Institute, Seattle, WA, USA (protocol IDs: IACUC00006 (K.J.M.); IACUC00108 (F.K.)).

## Figure Supplements

**Figure 1 – figure supplement 1:**
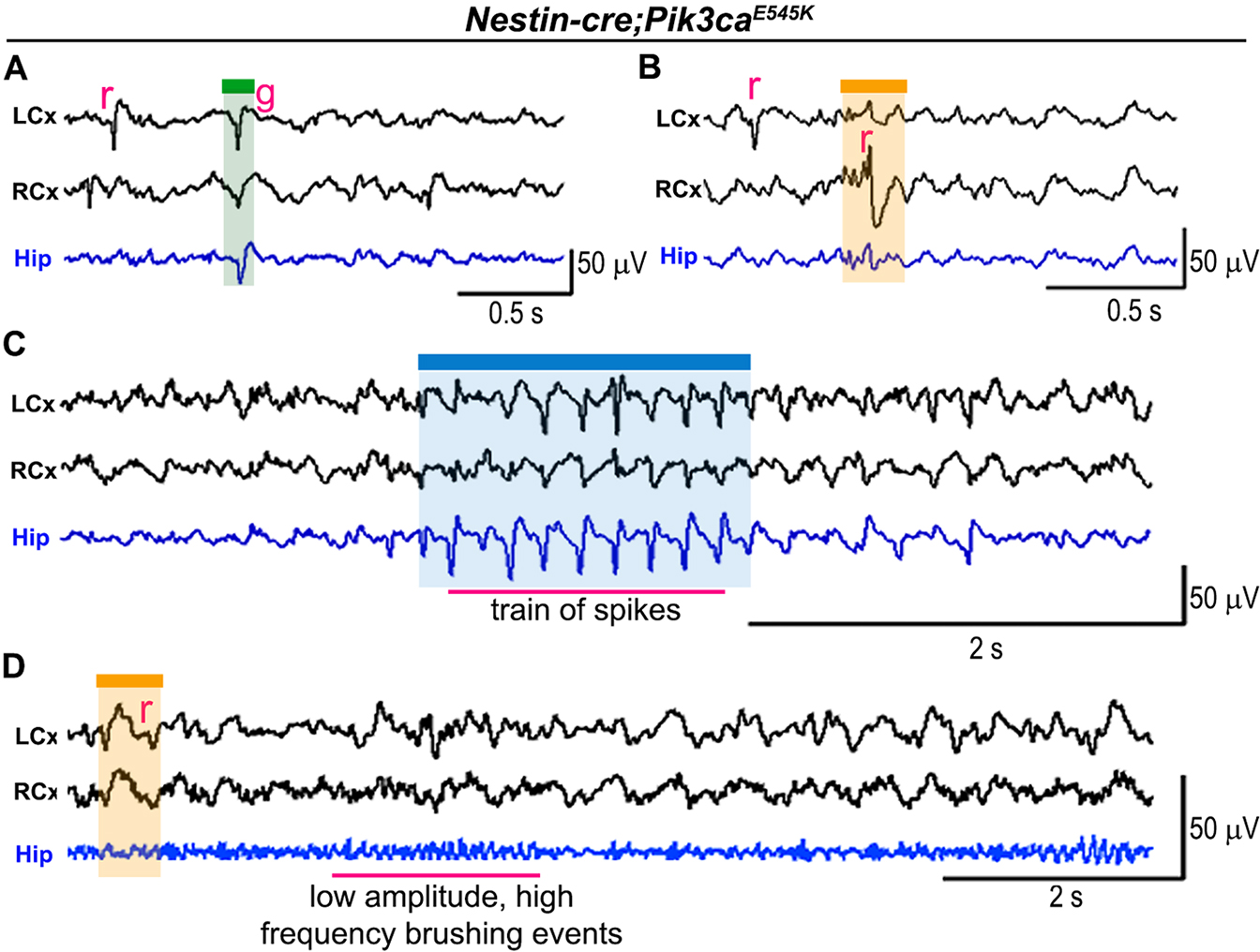
ECoG-LFP traces show different types of epileptiform activity in *Nestin-cre;Pik3ca^E545K^* brains. (**A,B**) Examples of different types of isolated spiking events in the neocortex and hippocampus taken from 2 different mutant mice showed a varied range of spike nature – in frequency, source of origin and amplitude. (**C**) Example of train of spiking events in the hippocampus and left cortex. (**D**) Example of low amplitude, high frequency brushing events. g, generalized; r, regional. Scalebars: 0.5s,50mV (A,B); 2s,50mV (C,D).

**Figure 2 – figure supplement 1:**
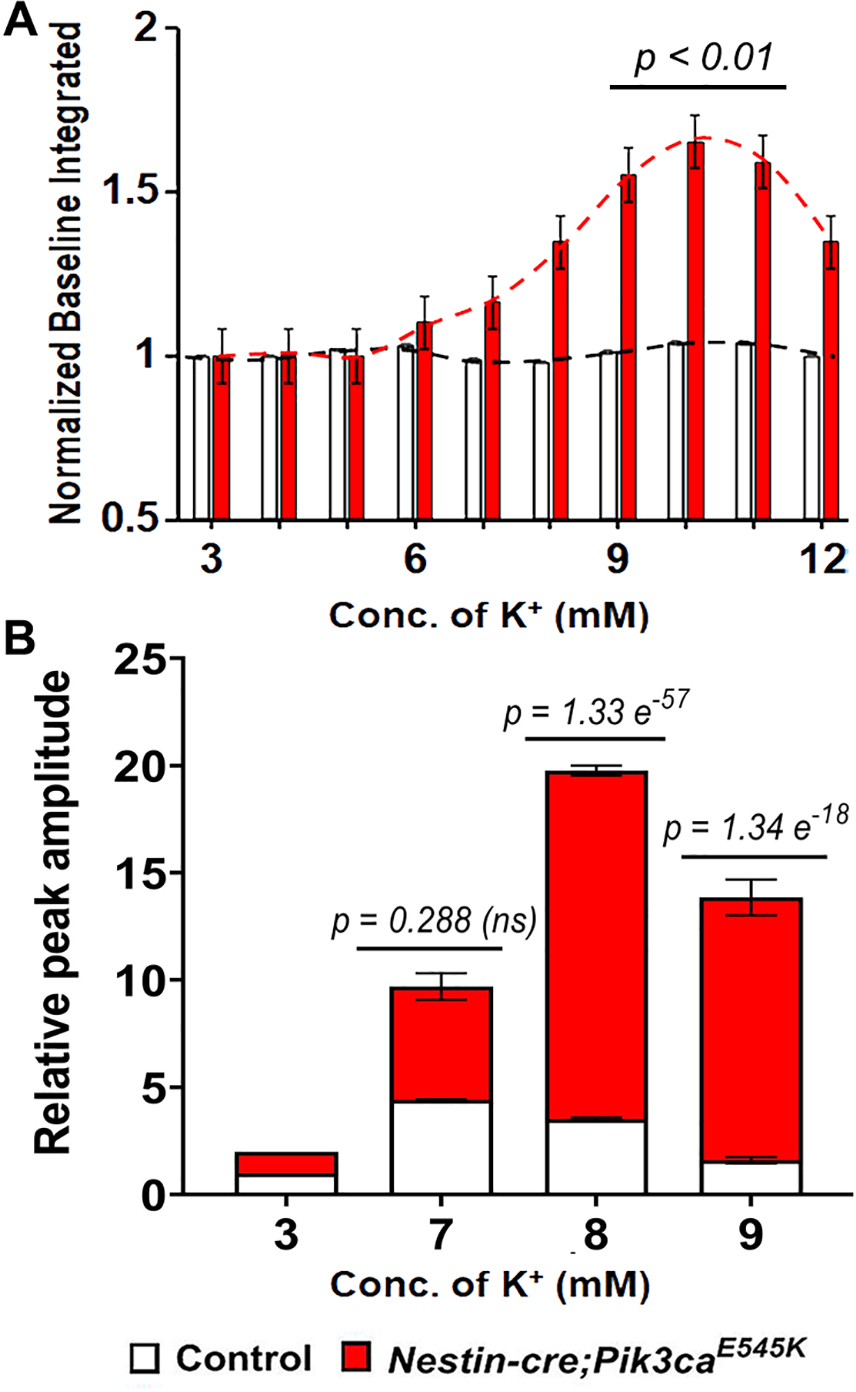
Enhanced neuronal network excitability in mutant CA1 across an extracellular potassium gradient. (**A**) Bar graphs depicted normalized baseline shift of integrated field potential traces in the mutant with increasing [K^+^] (refer to Figure 2). The mutant baseline was significantly higher than that of control especially in the 9-11mM range (t=3.343, degrees of freedom (df)=9). (**B**) Relative peak amplitude of mutant spiking was significantly higher than controls at higher potassium concentrations (F=266.1; df=564). Data is represented as mean ± SEM bar graphs; differences were considered significant at p<0.05; ns, not significant.

**Figure 3 – figure supplement 1:**
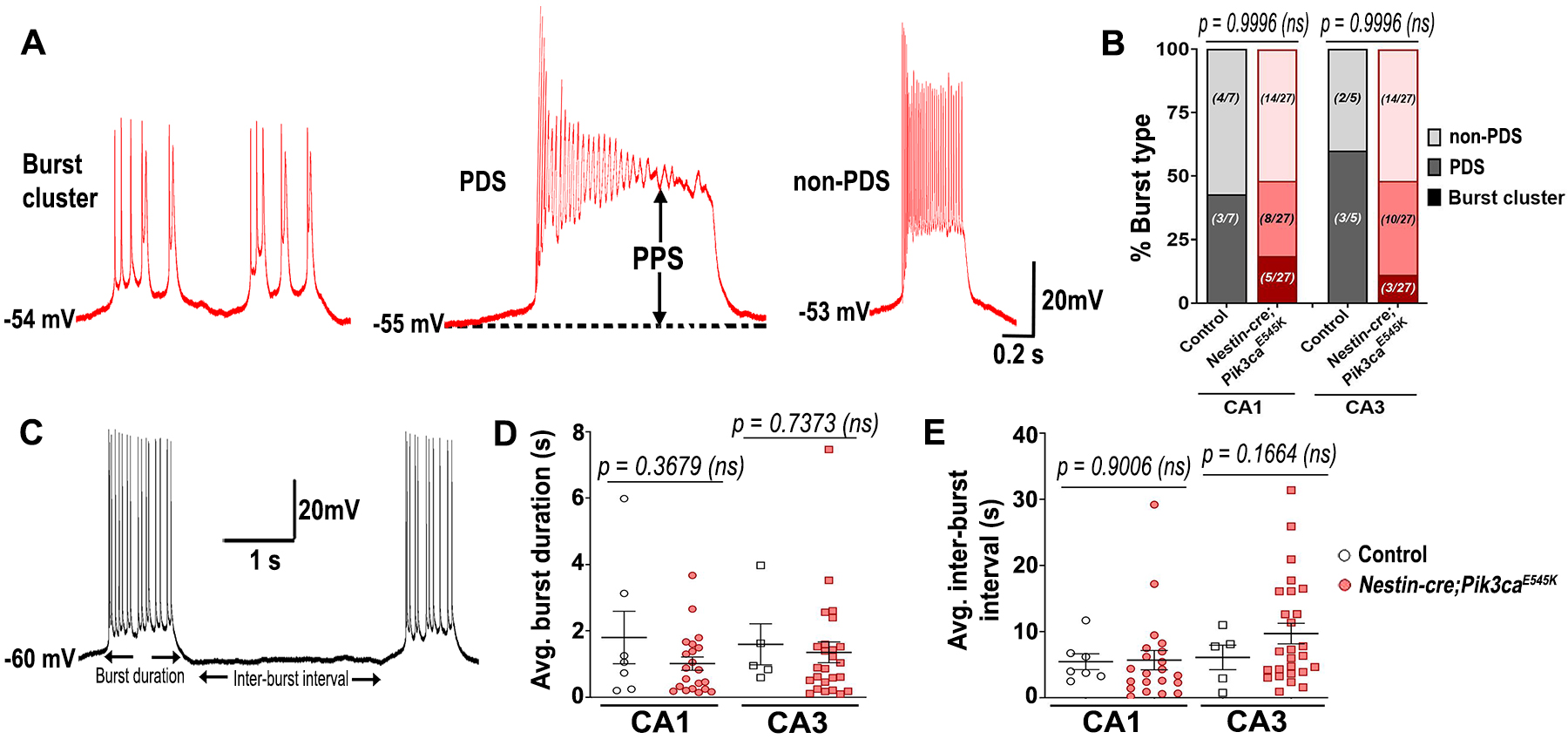
Spontaneous burst-firing features of control and mutant pyramidal neurons. (**A**) Representative traces for subtypes of burst firing, namely burst cluster, paroxysmal depolarization shift (PDS) and non-PDS plateau bursts. (**B**) Proportion of burst subcategories were not overtly different in control and mutant CA1 and CA3 regions (F= 7.355e^-12^; df=6); burst clusters were only seen in mutant cells. (**C**) Representative trace demonstrates how burst duration and inter-burst interval were calculated. (**D,E**) In both CA1 and CA3, average (avg.) burst duration (CA1: t=0.9644, df=6.793; CA3: t=0.3508, df=6.229) and inter-burst interval (CA1: t=0.1263, df=22.31; CA3: t=1.485, df=10.73) were not significantly different between control and mutant neurons. Data is represented as mean ± SEM scatter plots; differences were considered significant at p<0.05; ns, not significant; PPS, plateau potential shift. Scalebars: 0.2s,20mV (a), 1s,20mV (c).

**Figure 3 – figure supplement 2:**
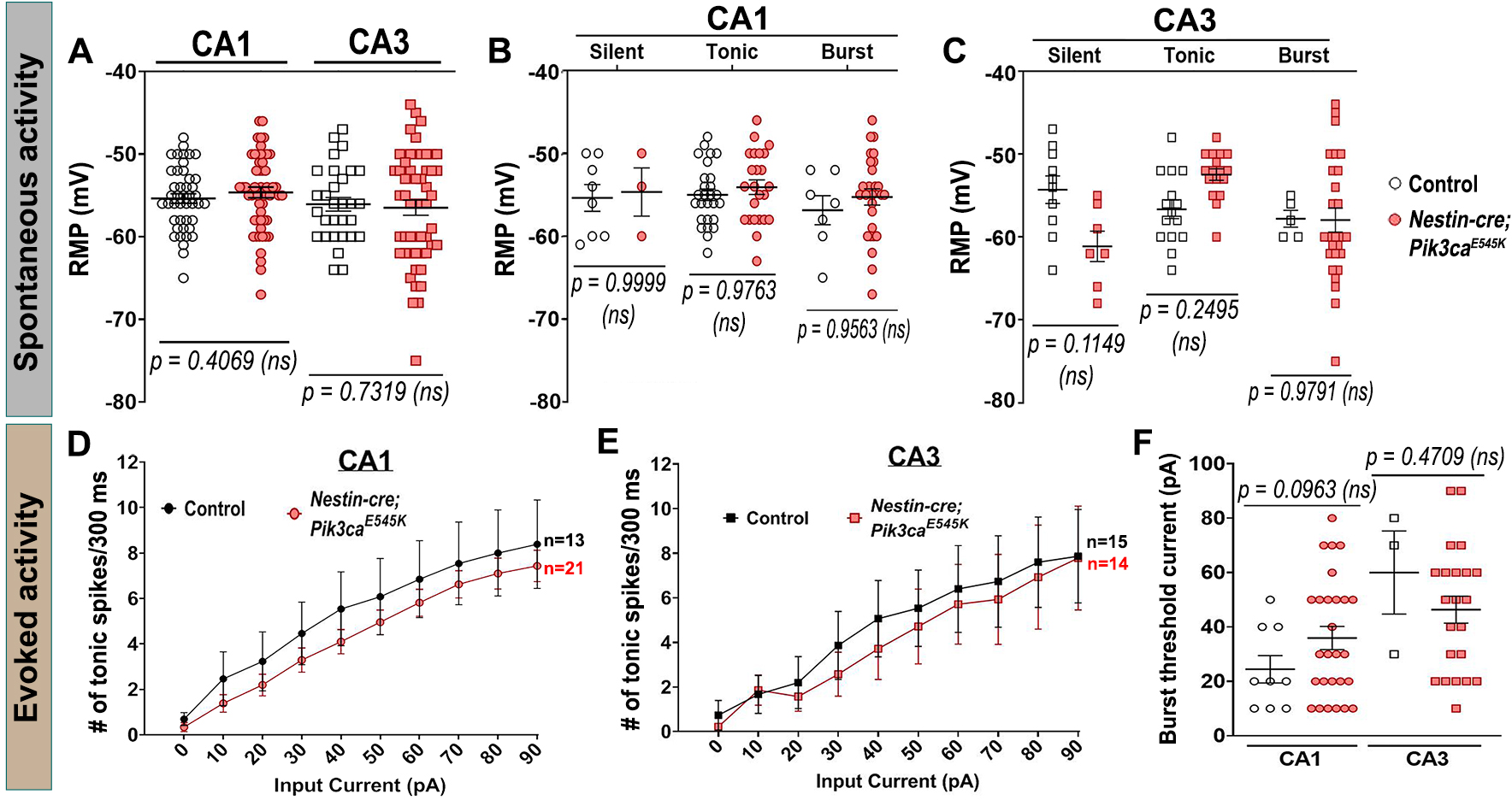
Resting membrane potentials and threshold currents for hippocampal neuronal firing remain unchanged by Pik3ca overactivation. (**A-C**) Resting membrane potential (RMP) was not significantly different between control and mutant neurons, both in CA1 (t=0.8331, df=94.99) and CA3 (t=0.3438, df=78.53), as well as among different firing types (CA1: F=0.9136, df=90; CA3: F=2.829, df=75). (**D,E**) No significant differences in evoked tonic spike frequencies were observed in the tested range of input depolarizing current (0-90pA), between control and mutant CA1 and CA3 neurons (CA1: F=13.07, df=320; CA3: F=4.844, df=270). (**F**) No overt differences in the current inducing the first burst were observed between control and mutant (CA1: t=1.744, df=20.29; CA3: t=0.8493, df=2.439). Data is represented as mean ± SEM scatter and line plots; differences were considered significant at p<0.05; ns, not significant.

**Figure 4 – figure supplement 1:**
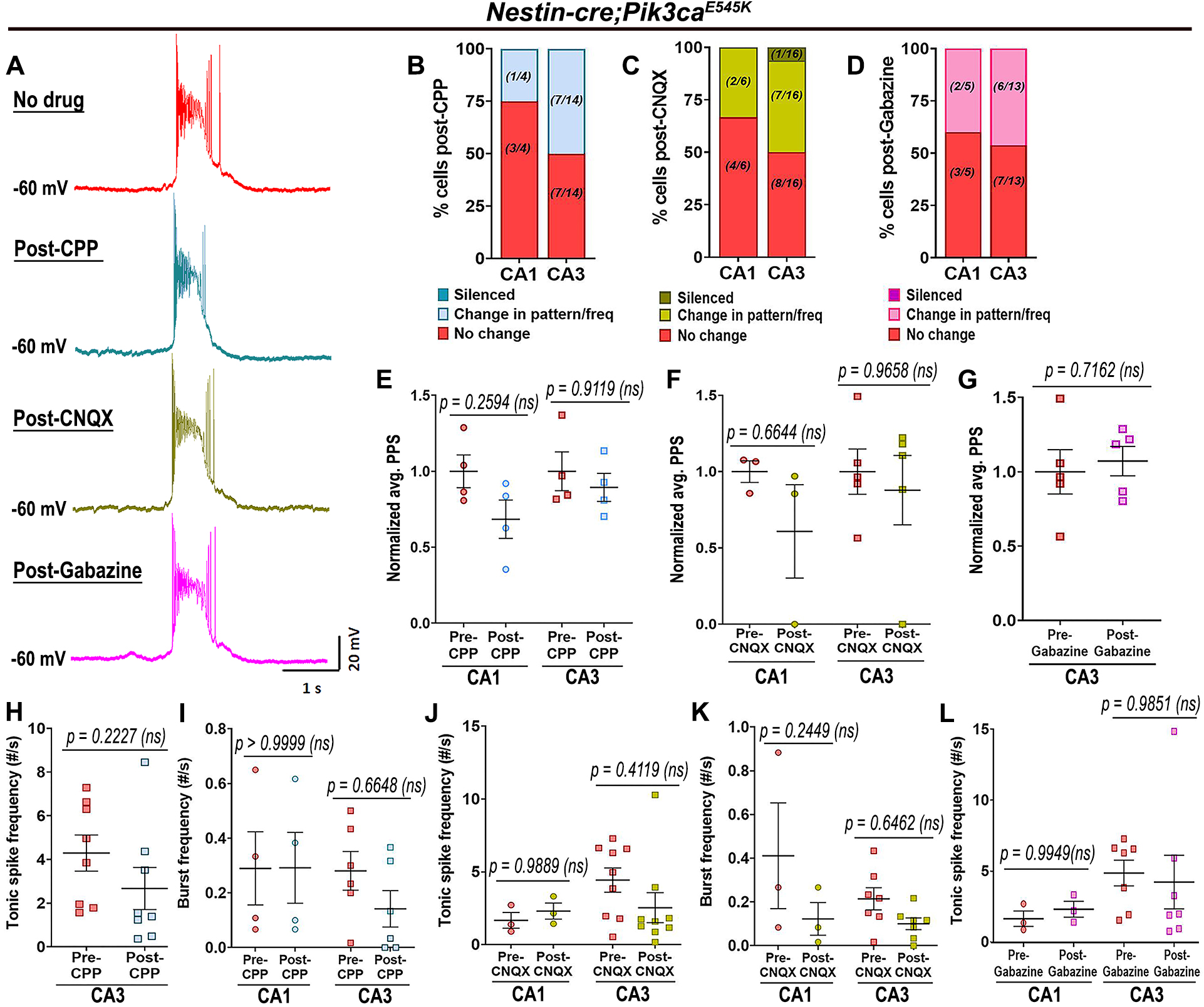
Pik3ca-related epileptiform activity is not dependent on glutamatergic or GABAergic inputs. (**A**) Representative spontaneous mutant trace and traces of the same cell post-treatment with glutamatergic receptor antagonists (CPP, CNQX) and with GABA antagonist gabazine showed no significant change in firing pattern. (**B-D**) Large proportion of mutant cells remained unaffected after being blocked from external glutamatergic or GABAergic inputs. (**E-L**) No significant differences in normalized plateau potential shift (PPS; CPP: F=3.391, df=12; CNQX: F=1.432, df=12; Gabazine: t=0.3904, df=4) or in tonic (CPP: t=1.876, df=7; CNQX: F=0.2766, df=20; Gabazine: F=0.0001130, df=16) and burst frequencies (CPP: F=0.5140, df=16; CNQX: F=5.218, df=16) were observed before and after administration of channel blockers. Data is represented as % bar graphs and mean ± SEM scatter plots; differences were considered significant at p<0.05; ns, not significant. Scalebar: 1s,20mV (A).

**Figure 5 – figure supplement 1:**
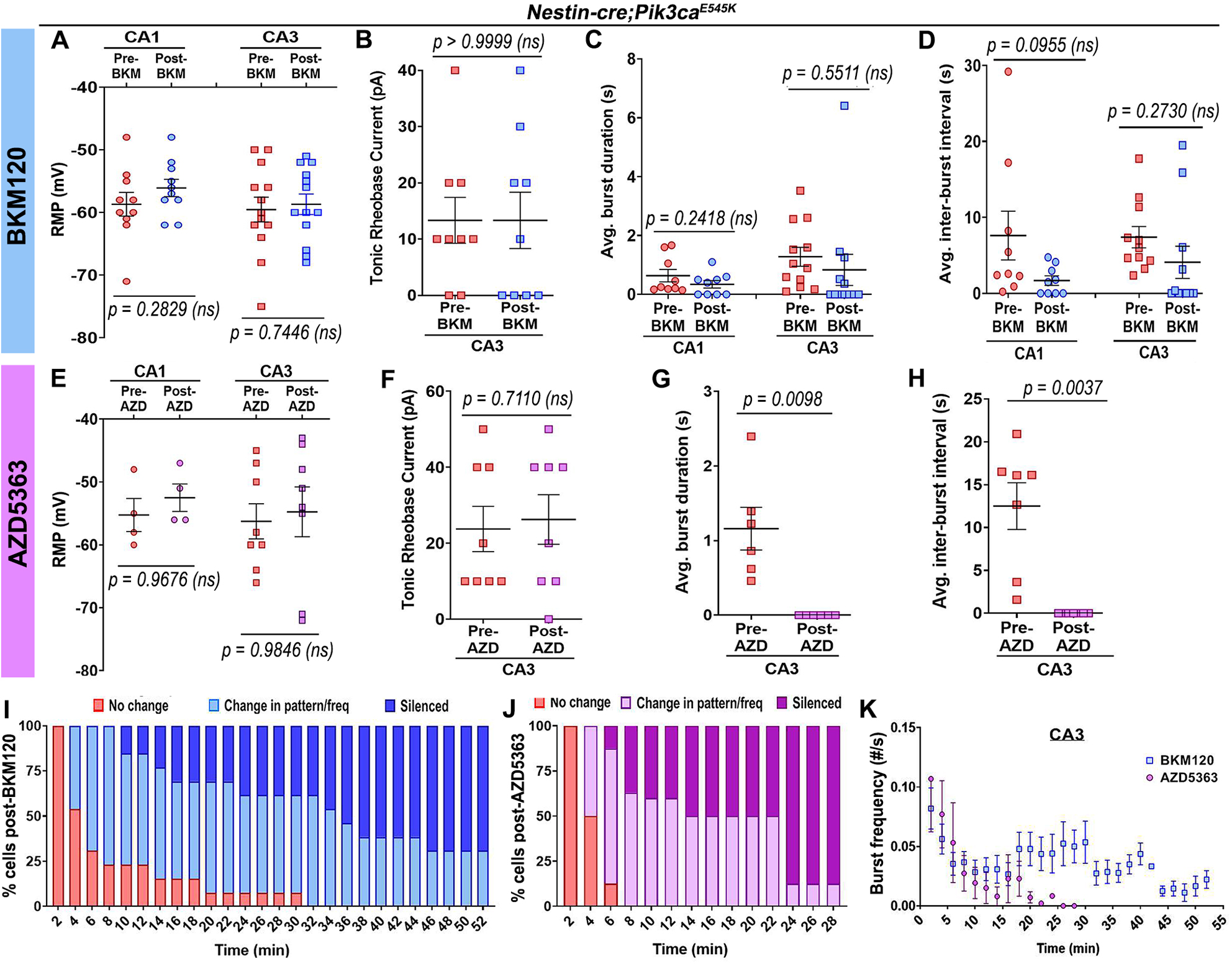
Acute treatment with BKM120 and AZD5363 has differential effect on *Nestin-cre;Pik3ca^E545K^* neuronal physiological properties. (**A-D**) Acute BKM120 treatment did not significantly affect RMP (F=0.9268, df=42), tonic rheobase current (t=0.000, df=8), average burst duration (CA1: t=1.264, df=8; CA3: t=0.6150, df=11) or inter-burst interval (CA1: t=1.889, df=8; CA3: t=1.160, df=10) in mutant hippocampal neurons. (**E-H**) Acute AKT inhibition by AZD5363 had no effect on the mutant RMP (F=0.3316, df=20) or rheobase current (t=0.3859, df=7), but significantly attenuated the average burst duration (t=4.052, df=5) and inter-burst interval (t=4.597, df=6), largely by suppressing bursts or silencing the cells. (**I,J**) Plots marked percentage of mutant CA3 “bursting” cells undergoing changes as a function of time, in response to acute administration of BKM120 (I) and AZD5363 (J). (**K**) Mutant plateau frequency significantly dropped as a function of time, where 0min marked the time of administration of BKM120/AZD5363 onto the brain slice. Data is represented as mean ± SEM scatter plots; differences were considered significant at p<0.05; ns, not significant.

**Figure 5 – figure supplement 2:**
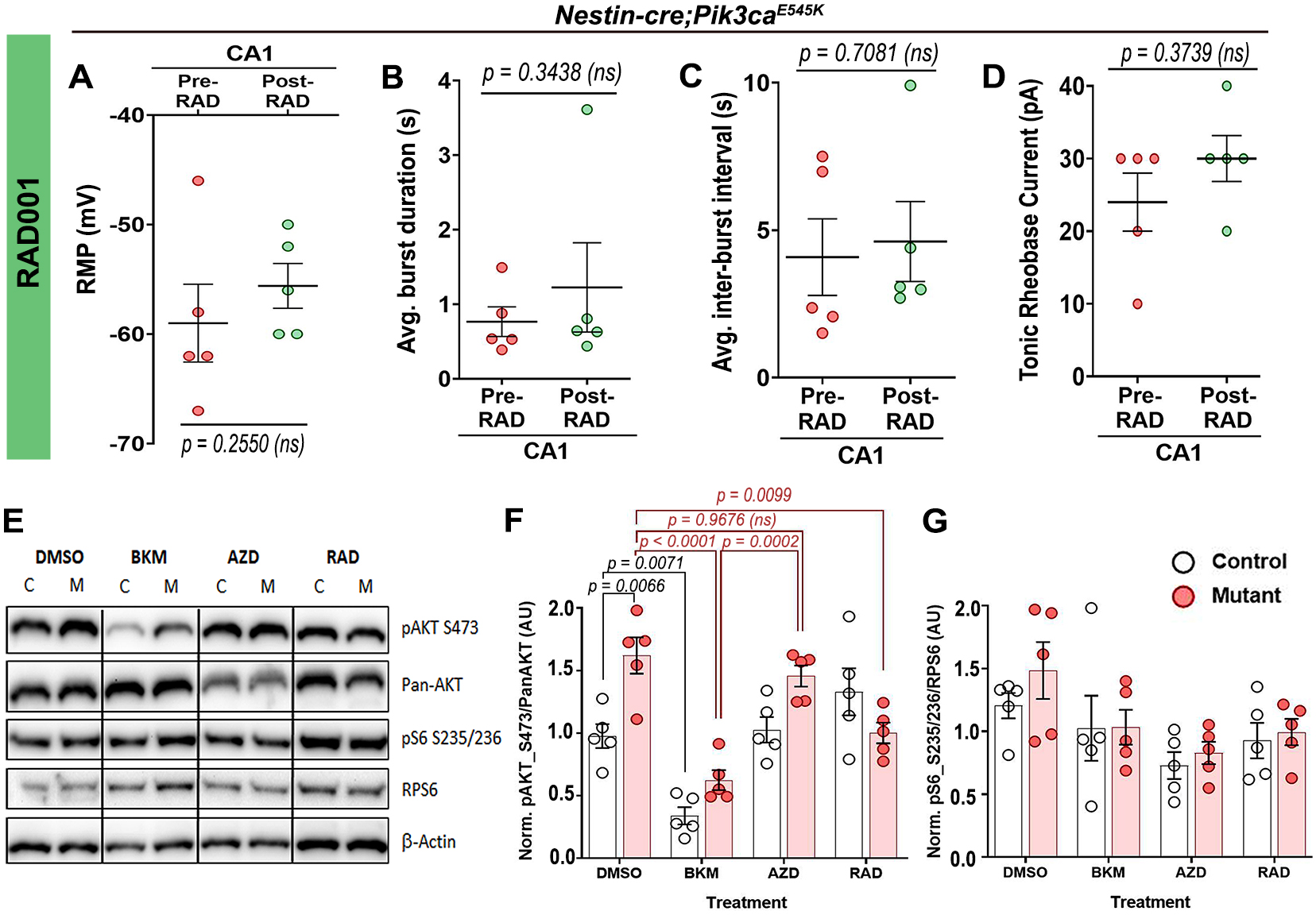
Acute treatment with RAD001 has no overt effect on mutant neuronal physiological properties. (**A-D**) Acute BKM120 treatment did not significantly affect the RMP (t=1.327, df=4), average burst duration (t=1.073, df=4), inter-burst interval (t=0.4023, df=4) or rheobase current (t=1.000, df=4) in mutant CA1 neurons. (**E**) Representative Western blots bands of pAKT_S473, pan-AKT, pS6-S235/236 and RPS6 (total S6) are demonstrated across different treatment (DMSO, BKM120, AZD5363, RAD001). (**F,G**) Quantifications of normalized ratios of phosphoproteins over total proteins showed significant reduction of pAKT_S473 in response to acute BKM120 and RAD001 treatment but not post-AZD5363 (F=22.98, df=32); pS6_S235/236 in mutant had a decreasing trend in response to all pathway drugs (F=4.485, df=32). Data is represented as mean ± SEM scatter plots and bar graphs; differences were considered significant at p<0.05; ns, not significant.

